# Trilobite moulting behaviour variability had little association with morphometry

**DOI:** 10.1101/2022.12.12.520015

**Authors:** Harriet B. Drage

**Affiliations:** Institute of Earth Sciences, University of Lausanne, 1015 Lausanne, Switzerland

**Keywords:** behaviour, moulting, morphometry, morphospace, multimodal distribution, trilobite

## Abstract

Trilobite moult assemblages preserved in the fossil record show high variability in moulting behaviour and their resulting moult configurations. The reasons for this variability, and the impacts it might have had on their evolutionary trajectories, are unknown and have rarely been investigated quantitatively. A large dataset of trilobite moult morphometric measurements is presented and statistically analysed for associations between moulting behaviour and morphometry. Results indicate little significant statistical association between the two; only between moulting behaviour (usually generalised moult configuration) and the variances and means of thoracic tergite number, thorax length, and pygidium width. Anterior cranidium width, cranidium length, cephalothoracic joint width, thorax width, pygidium length, and total body length all have non-significant associations with moulting behaviour. Moult specimens showing inversion of the librigenae generally have more thoracic tergites, a correspondingly longer thorax, and a narrower pygidium. Thoracic tergite count and pygidium measurements may have multimodal distributions. Principal Components Analyses and Non-Metric Multidimensional Scaling analyses suggest minor differences in the extent of morphometric variation for specimens showing different moulting behaviours, but little difference in the region of morphospace they occupy. This may indicate that trilobite species using Salter’s mode of moulting had more constrained morphologies, potentially related to facial suture fusion in some groups. Overall, these results do not suggest a strong association between moulting behaviour variation and morphometry in trilobites, leaving open for further study the mystery of why trilobites were so variable in their moulting, and whether this contributed to their long evolutionary reign or ultimate extinction.

**PLAIN LANGUAGE SUMMARY:** Trilobites were an important and globally abundant group of arthropods (animals with an exoskeleton and jointed limbs) that lived ~521-251 million years ago. The exoskeletons of arthropods are crucial because they provide protection against predators and parasites, but also restrict their growth. All living and extinct arthropods must therefore periodically moult (shed) their exoskeletons; an incredibly risky event during which many individuals die. Due to its importance, it is presumed that exoskeleton moulting impacted the broad-scale evolution of arthropod morphology (their physical characteristics), behaviour, and ecology. Trilobite moults are preserved in great number in the fossil record, and this can tell us much about their moulting behaviour. Additionally, trilobites appear to be unique in showing many different moulting behaviours. However, we do not know why trilobites were so variable in their moulting behaviour, or what impact this had on their evolution. In this study, a large dataset of trilobite moulting behaviours and their body proportion (morphometry) measurements is presented and analysed to answer: ‘Was variability in trilobite moulting behaviour related to differences in their morphometry?’ The results suggest that there was little association between the moulting behaviours shown by trilobites and their morphometry. Species showing the different moulting behaviours had overall similar morphologies, although for one moulting behaviour this seemed more limited. Only thorax length and segmentation (the central part of the body), and pygidium (‘tail’) width, significantly differed between species showing the different moulting behaviours. This study does not indicate a strong relationship between moulting behaviour and morphology in trilobites. This is unexpected, and leaves open the mystery of trilobite moulting variability.

## INTRODUCTION

Exoskeleton moulting is a key recurrent event in the life histories of all euarthropods. The exoskeleton protects individuals from predation and parasitism (Ewer, 2005), but is naturally restrictive so must be periodically moulted for significant growth and developmental changes to occur. This moulting is energetically expensive and leaves the individual periodically more vulnerable to predation during exuviation (exiting of the old exoskeleton) and during reinforcement of the new exoskeleton (Vevea and Hall, 1984). The behaviours and mechanisms involved in moulting are therefore inherently linked to euarthropod morphology, development, ecology, and broad-scale evolution (Brandt, 2002; Daley and Drage, 2016). The fossil record of euarthropod moulting is extensive (Daley and Drage, 2016), allowing exploration of the interactions between moulting behaviour and these key biological facets.

Trilobites have a rich fossil record of moults with which to explore the impact of moulting behaviours on their evolution, owing to their exoskeletons reinforced with calcite that have a high preservation potential (Teigler and Towe, 1975). Further, trilobites are unusual because they appear to show a greater range of moulting behaviours and preserved moulting configurations (the configuration of disarticulated sclerites preserved in the fossil record and reflecting moulting movements) than other euarthropod groups with apparently specialised moulting behaviours leading to recognisable configurations (Brandt, 2002; Daley and Drage, 2016; Drage, 2019a,b). Not only do trilobites show high interspecific variability in their moulting, but also seemingly extensive intraspecific variation, with different individuals of the same species preserving varied configurations (see Drage et al., 2018a). Trilobites are also one of the most successful groups of euarthropods based on their diversity of ecological niches, abundance, global distribution, and evolutionary timespan (Fortey and Whittington, 1989; Fortey and Owens, 1999; Webster, 2007).

Trilobites show the evolution of a variety of sophisticated sutures involved in moulting, for example, the cephalic sutures, which may exist solely to facilitate moulting through disarticulating the librigenae and producing an anterior exuvial gape (Stubblefield, 1959; Hughes, 2007; Hou et al., 2017; Du et al., 2019). The facial sutures were also secondarily lost in some groups (see Daley and Drage, 2016) or occasionally fused during development (Drage et al., 2018b). A variety of other cephalic sutures were used for moulting in some groups, such as an anterior marginal suture, rostral and hypostomal sutures, and ventral median sutures, as well as important use of the cephalothoracic joint for moulting (Henningsmoen, 1975; Fortey, 2001; Budil and Bruthansova, 2005; Drage, 2019b). Consequently, trilobites display several modes of moulting, which can be summarised into (see Table 1):

1. use of the cephalic exuvial sutures (various combinations of facial, rostral, and hypostomal sutures depending on morphology), termed the Sutural Gape mode of moulting
2. use of the cephalothoracic joint for moulting, termed Salter’s mode of moulting
3. use of the marginal suture along the anterior cephalic margin (Henningsmoen, 1975; Drage, 2019b).

**TABLE 1.** Moulting behaviour terminology used throughout this work. Details of the specific moult configurations described for trilobites can be found in Drage et al. (2018a).

|  |  |  |
| --- | --- | --- |
| **Modes of moulting used in this study** | **Sutural Gape mode of moulting** | Mode of moulting that involves only opening of the cephalic sutures, which may comprise the facial, rostral, and/or hypostomal sutures. |
|  | **Salter's mode of moulting** | Mode of moulting that involves only opening of the cephalothoracic joint during moulting, causing disarticulation of the cephalon. |
|  | **Hybrid mode of moulting** | Mode of moulting combining both the Sutural Gape and Salter's modes, with both cephalic sutures and the cephalothoracic joint opening during moulting. |
| **Generalised moult configurations used in this study** | **Cephalic sutures configuration** | Preserved moult configurations that have variously opened cephalic sutures, and so have related cephalic sclerites disarticulated from the body (librigenae, rostral plate, and/or hypostome). This includes moults in Harrington's, Nutcracker, and Axial Shield configurations (of Drage et al., 2018a). |
|  | **Cephalic sutures + inversion configuration** | As above, but with some of the disarticulated cephalic sclerites, usually librigenae, either horizontally or vertically flipped, as in the Somersault and McNamara's configurations respectively (Drage et al., 2018a). |
|  | **Salter's configuration** | Moult configuration with the entire cephalon disarticulated at the cephalothoracic joint, and horizontally inverted compared to the remaining thoracopygon (Drage et al., 2018a). |
|  | **Zombie configuration** | Moult configuration with the entire cephalon disarticulated at the cephalothoracic joint, and displaced from the remaining thoracopygon (Drage, 2019b). |
|  | **Henningsmoen's configuration** | Moult configuration resulting from the hybrid mode of moulting, with the cephalon disarticulated at the cephalothoracic joint, but also the librigenae, rostral plate, and/or hypostome disarticulated (Drage et al., 2018a). |

In concert with the individual movements of trilobites, these produce an array of moult configurations that are found preserved in the fossil record (Figure 1). Drage et al. (2018a) described and named a number of these pertaining to the Sutural Gape and Salter’s modes of moulting using an exceptionally preserved, diverse sample of *Estaingia bilobata*.

**FIGURE 1.**
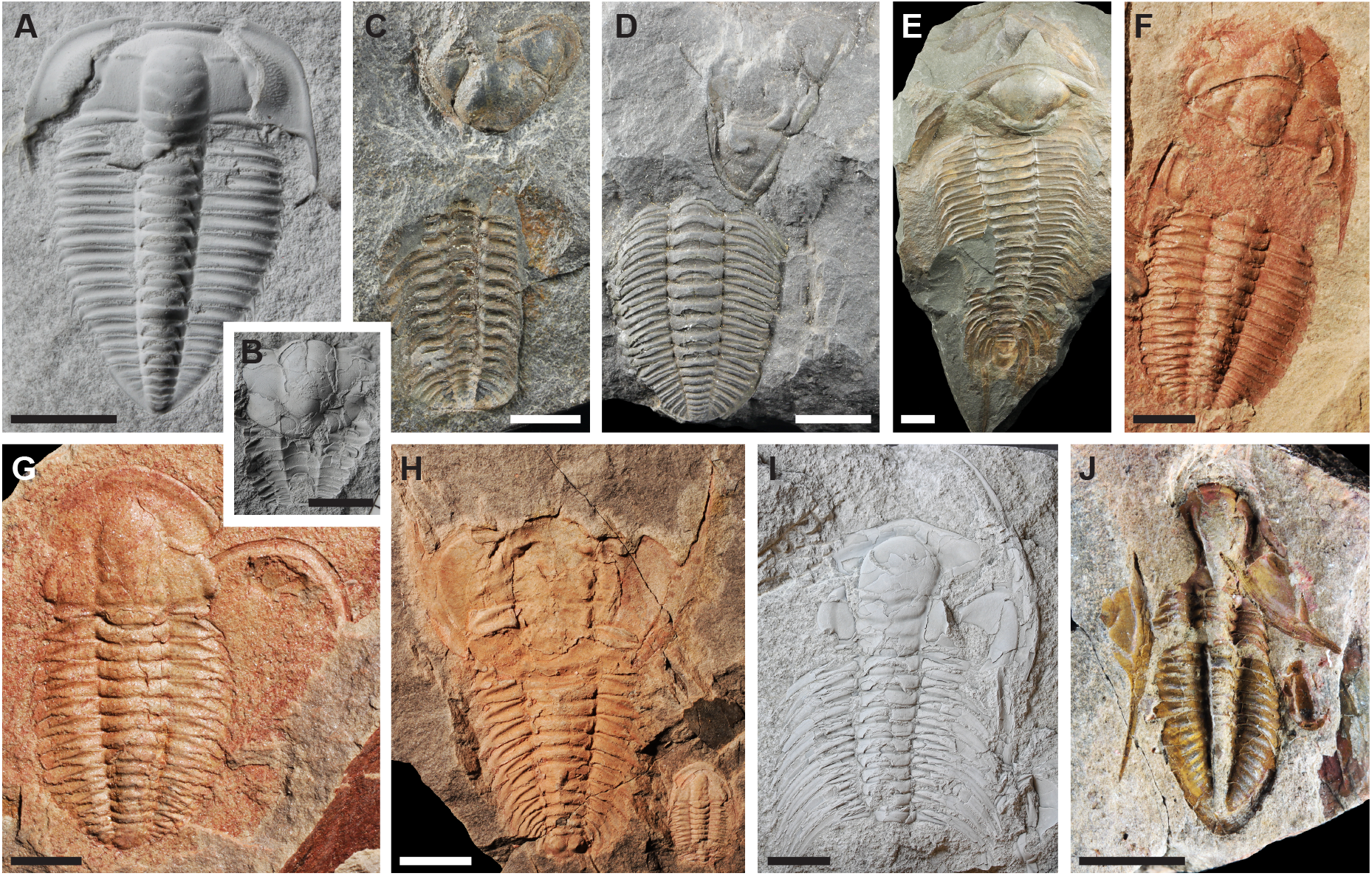
Trilobite moult configurations featuring in the dataset presented here. Moults representing the following generalised moult configuration groups used in this study: cephalic sutures configuration group; cephalic sutures + inversion configuration group; Salter’s configuration; Zombie configuration; Henningsmoen’s configuration. A: *Olenus truncatus*, PMU unnumbered specimen, B: *Ellipsocephalus* sp., PMU 28642, C: *Trimerocephalus mastophthalmus*, NHMUK I.5100, D: *Ceraurus pleurexanthemus*, NHMUK 42341, E: *Paradoxides bohemicus*, NHMUK 42440, F: *Estaingia bilobata*, SAM-P 46956, G: *Estaingia bilobata*, SAM-P 43767, H: *Redlichia takooensis*, SAM-P 43593, I: *Accadoparadoxides pinus*, PMU 25995, J: *Phillipsia* sp., NHMUK I.1092. Scale bars = 5 mm for A, B, F, G, J; 10 mm for C-E, H, I.

Several early works emphasised the potential important impacts of trilobite moulting behaviour variability on their evolutionary trajectories (Henningsmoen, 1975; Whittington, 1990; Budil and Bruthansova, 2005) and the potential impact of this behaviour on their extensive success and ultimate decline (Brandt, 2002). Recent studies have described moulting in specific trilobite groups, such as Zong (2020) on moults of *Ovalocephalus tetrasulcatus* arranged end-to-end, synchronised moulting behaviour of several oryctocephalid species preserving diverse moulting configurations by Corrales-García et al. (2020), and ontogenetic moulting sequence trends in *Arthricocephalites xinzhaiheensis* by Wang et al. (2021). These studies have further increased support for inter- and intraspecific variability in moulting across Trilobita and its links with other aspects of morphology and behaviour. Indeed, all descriptions of the fossil record of moulting serve to further draw attention to and increase our understanding of this crucial behaviour in Euarthropoda. Drage (2019a) presented the first broad-scale study quantifying the extent of inter- and intraspecific variability in trilobite moulting, also exploring potential trends in moulting with taxonomic assignment and geological history. However, still little is known about why this moulting variability existed, the impacts it may have had on the evolutionary history of the group, and how moulting interacted with morphology.

This study uses the large dataset of interspecific trilobite moulting variability presented by Drage (2019a) and corresponding moult morphometric (comparative body proportion) measurements to test for an association between moulting behaviour and morphology. Variance in morphometry may have impacted individual moulting behaviours, for example, differences in cephalic proportions might have changed exuvial gape sizes, making some behaviours more complex or disarticulations more likely. Similarly, moulting may have itself imposed constraint on trilobite morphological evolution, making certain moulting behaviours less risky. This work therefore aims to determine whether there is a significant association between the unusual interspecific moulting variability observed in trilobites and their morphometry.

## METHODS AND MATERIALS

### Data collection

The dataset analysed here consists of descriptions of moulting configurations for 617 individuals of 238 trilobite species, which was originally presented by Drage (2019a). See the methodology described in Drage (2019a) for further details of specimen choice, location, and recording. The majority of specimens in the dataset were described and measured in person (by H.B. Drage). However, 46 out of 617 moults were included based on photographs from the descriptive literature (see Supplementary data); these were measured using Fiji (Schindelin et al., 2012).

Extensive literature searches were performed to confirm metadata regarding geological age range, taxonomic assignment, developmental stage, and species thoracic tergite count. However, some species descriptions are not readily available in the accessible literature; for example, information on complete thoracopygidia is not available for trilobite species originally described from isolated cephala (Whittington et al., 1997). Specimens that might have been of a late juvenile developmental stage, based on thoracic tergite number and relative size, were removed from the dataset. This is because the inclusion of juveniles could bias results as immature individuals have been shown to moult differently to their adult counterparts in some species (Drage et al., 2018b; Wang et al., 2021), and ontogenetic studies show that morphology can differ extensively between developmental stages (Park and Choi, 2011). It can be difficult to distinguish late-stage meraspides, which can have a full complement of thoracic tergites, from holaspides, as many trilobite species are not known from a sufficient number of juvenile specimens to describe their ontogenies, and some species also had variable numbers of tergites during functional adulthood (Hughes, 2007; Park and Choi, 2011). Thus, all species in the dataset have been checked in the descriptive literature to confirm adulthood, if possible, but a level of error resulting from accidental inclusion of late-stage meraspides remains plausible. In addition, specimens for which a moult assignment was uncertain were removed from the dataset, such as when there was too little associated material on the rock sample (only isolated thoracopygidia or cephala present) or material was highly fragmented (see moult designation criteria in Daley and Drage, 2016).

Linear measurements of key body dimensions were taken using digital callipers at the millimetre scale for each of the moult specimens, where possible given preservation, during the same period of data collection as for the data presented in Drage (2019a). These included: anterior cranidium width (tr., between the anterior-most dorsal extension of the facial sutures, if present); cranidium axial length (sag.); cephalothoracic joint width (tr.); thorax maximum width (tr., without pleural spines); thorax axial length (sag.); pygidium maximum width (tr., without pygidial spines); pygidium axial length (sag.); and total axial body length (sag.). The thoracic tergite number was also recorded.

Drage et al. (2018a) named and figured a diversity of moult configurations, however, it was not feasible to assign these named configurations to the moult specimens in this dataset as many specimens display non-exceptional preservation or natural variability in sclerite placement, so do not fit the ideal descriptions given by Drage et al. (2018a). For example, if fossilisation is insufficiently efficacious to preserve the underlying integument, Harrington’s configuration would be preserved as the Nutcracker configuration, with all disarticulated cephalic sclerites separated (following Drage et al., 2018a). Accordingly, moults that may have been preserved in any of the Nutcracker, Harrington’s, or Axial Shield configurations (of Drage et al., 2018a) were grouped together in the dataset as ‘cephalic sutures’ configurations, which is defined as moults with the cephalic exuvial sutures open but does not distinguish between the placement of these in the preserved moult. Similarly, McNamara’s and Somersault configurations (of Drage et al., 2018a) were combined as ‘cephalic sutures + inversion’ configurations, defined by both opening of the cephalic exuvial sutures and horizontal or vertical inversion of the librigenae during moulting. Ultimately, each moult specimen was assigned both a mode of moulting (Sutural Gape, Salter’s, or hybrid) and a generalised moult configuration (cephalic sutures, cephalic sutures + inversion, Salter’s, Zombie, or Henningsmoen’s configuration); see full descriptions in Drage et al. (2018a) and Table 1. The ‘hybrid’ mode of moulting includes only specimens preserved in Henningsmoen’s configuration, because this is a hybrid configuration displaying opening of both the cephalic sutures (Sutural Gape mode) and cephalothoracic joint (Salter’s mode) for moulting (Drage et al., 2018a).

### Dataset composition

Table 2 gives the number of moult specimens included in the dataset for each mode of moulting and generalised configuration. Specimens spanned most currently named trilobite orders, though extensive high-level taxonomic uncertainty remains for the group (e.g., the suggested paraphyly of Ptychopariida; Adrain, 2011). All raw data is freely available in the associated Supplementary data file.

**TABLE 2.**
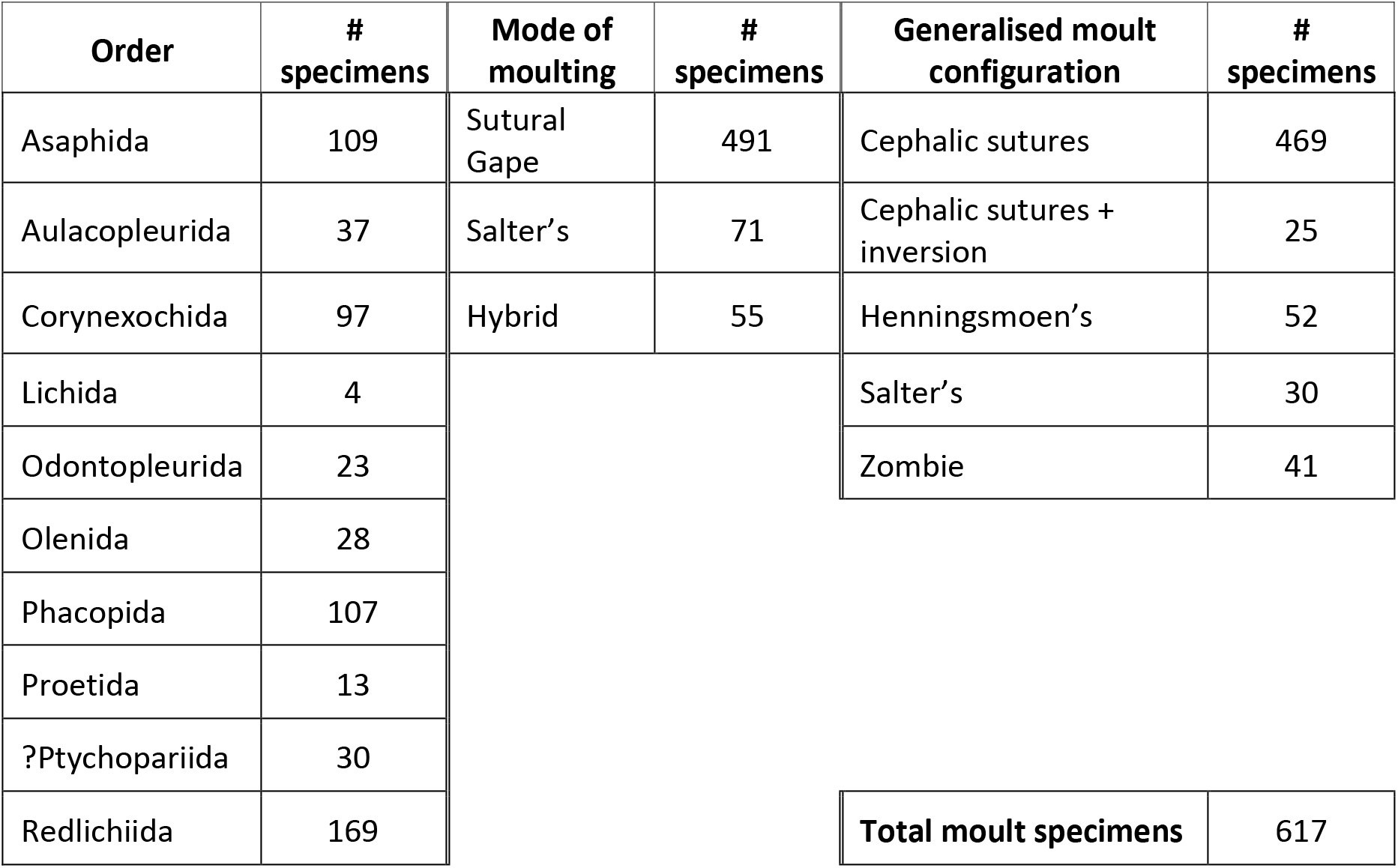
Number of specimens included in the dataset for each trilobite order, based on taxonomy of Adrain (2011). Count of specimens for each mode of moulting and generalised moult configuration (see Table 1 definitions) included in this study. Raw specimen data available in the Supplementary data file.

### Analyses

To test for associations between trilobite moulting behaviour and morphometry several targeted statistical analyses were performed. All analyses were performed and plots produced in R (Core Team, 2021) using RStudio (RStudioTeam, 2020) and the following additional packages: car (Fox and Weisberg, 2019); ColorBrewer (Brewer et al., 2003); ggplot2 (Wickham, 2016); ggpubr (Kassambara, 2020); jmv (Selker et al., 2021); mixtools (Benaglia et al., 2009); MorphoTools2 (Koutecky, 2015; Šlenker et al., 2022); stats (Core Team, 2021); tidyverse (Wickham et al., 2019); vegan (Oksanen et al., 2020). All analyses were performed on morphometric data of specimens grouped into both mode of moulting and generalised moult configuration (see Table 1). Descriptive statistics were produced for all morphometric variables.

Geometric morphometric analyses were performed to determine whether specimens showing the same moulting behaviours grouped together in morphospace. These consisted of both Principal Components Analyses (PCA) and Non-Metric Multidimensional Scaling (NMDS) analyses. PCA is traditionally used for biological morphometric analyses but cannot include data entries that contain missing information as it uses a Euclidean distance metric (Koutecky, 2015); this is unfortunately the case for this dataset, and frequently for fossil data. NMDS analysis uses alternative distance metrics (Buttigieg and Ramette, 2014), and represents a non-parametric alternative to PCA that can often be suitable for biological data due to their frequent violation of normality; these were therefore performed with all data included as a comparison to PCA. However, NMDS is not ideal in isolation because analyses can become stuck within local dataset minima, and it is not as informative as PCA because it cannot provide a breakdown of the influence of the original variables on the results. NMDS analyses produced Euclidean stress values of 0.12-0.13, which is considered suitable for reliable inference (Buttigieg and Ramette, 2014). NMDS analyses included all 617 specimens, while for PCA the total body length and pygidium length and width variables were removed due to their high proportions of incomplete data. With subsequent removal of incomplete data entries, this left 273 specimens for PCA. An eigenvector plot was produced to show the influence of the original measurement variables on the PCA morphospace. Ellipses on the PCA and NMDS plots represent the regions where new independent observations will fall with a 0.95 probability level. Following NMDS, non-parametric ANOSIM analyses were performed to test whether the dissimilarity in morphometry between each moulting group was greater than within each group (i.e., whether the moulting groupings would naturally group together based on morphometry; Buttigieg and Ramette, 2014).

Differences in average thoracic tergite number and measurement variable with moulting behaviour were tested for using Kruskal-Wallis chi-squared analyses. Post-hoc Wilcoxon pairwise analyses were performed when a significant Kruskal-Wallis result was obtained. Multivariate Analysis of Covariance (MANCOVA) tests were used to explore the impact moulting behaviour had (the independent variable) on each morphometry variable (the dependent variable), while controlling for all other variables (the covariates). Controlling for all covariates simultaneously removed the impact of allometric body scaling and related morphological change on each MANCOVA test, so these potentially confounding and interdependent variables do not impact the result. Scatter plots with linear regressions predicted by the MANCOVA models were produced to visualise any differences between the moulting groups. A significance threshold of *α* = 0.05 was used throughout the analyses, though all relevant *p* values are reported for interrogation.

The distributions of data present were explored through qualitative interpretation of the linear regression plots and morphospace plots previously mentioned, followed by production of histograms for each variable. The fit of each dataset variable to a single distribution or a multiple mixed set of overlapping distributions was analysed using an iterative expectation-maximisation (EM) algorithm method, and its resulting distribution models and log-likelihood values (Benaglia et al., 2009). The less negative the log-likelihood figure at convergence of the estimate, the better the distribution model from the EM algorithm fits the data.

## RESULTS

### Dataset moulting variability is high

As fully detailed in Drage (2019a), interspecific moulting variability in the dataset is high. However, most specimens are preserved as the cephalic sutures configuration group (469 specimens). Inversions of exoskeleton sclerites (including Salter’s and cephalic sutures + inversion configurations) are generally less common in the dataset (55 specimens total). Moults manifesting as hybrids between the Sutural Gape and Salter’s modes of moulting (the hybrid mode) were similarly present (55 specimens), likely due to differences in degradation and breakage of the integument during moulting, and preservational impacts like minor transport of individual sclerites by water currents in non-exceptional preservational environments.

### Moulting behaviours are not distinguished by morphospace occupation

Variance in PC1 of the PCA plots (Figure 2A-C) can be explained by differences in anterior cranidium width (tr.), cranidium length (sag.), cephalothoracic joint width (tr.), thorax width (tr.), and partially thorax length (sag.); that is, all included continuous measurement variables. Variance in PC2 (Figure 2A-C) can be explained almost entirely by thorax tergite number, and partially by thorax length (sag.). Dataset variation is represented by PC1 at 73% and PC2 at 20% (Figure 2A, B). The PCA plots suggest no difference in the region of morphospace occupied by the different moulting modes (Figure 2A) or generalised moult configurations (Figure 2B), as the ellipses are nested within each other. The NMDS plots support this finding, with the moulting mode (Figure 3A) and configuration (Figure 3B) group ellipses also being entirely nested. These results show that the different moulting mode and configuration groups cannot be distinguished based on their overall morphometries. ANOSIM tests support this, being non-significant for both moulting mode (*R* =-0.0541, *p* = 0.856) and configuration (*R* =-0.0632, *p* = 0.928), supporting the nesting of ellipses in the NMDS analyses and confirming that there no significant difference between the moult groupings.

**FIGURE 2.**
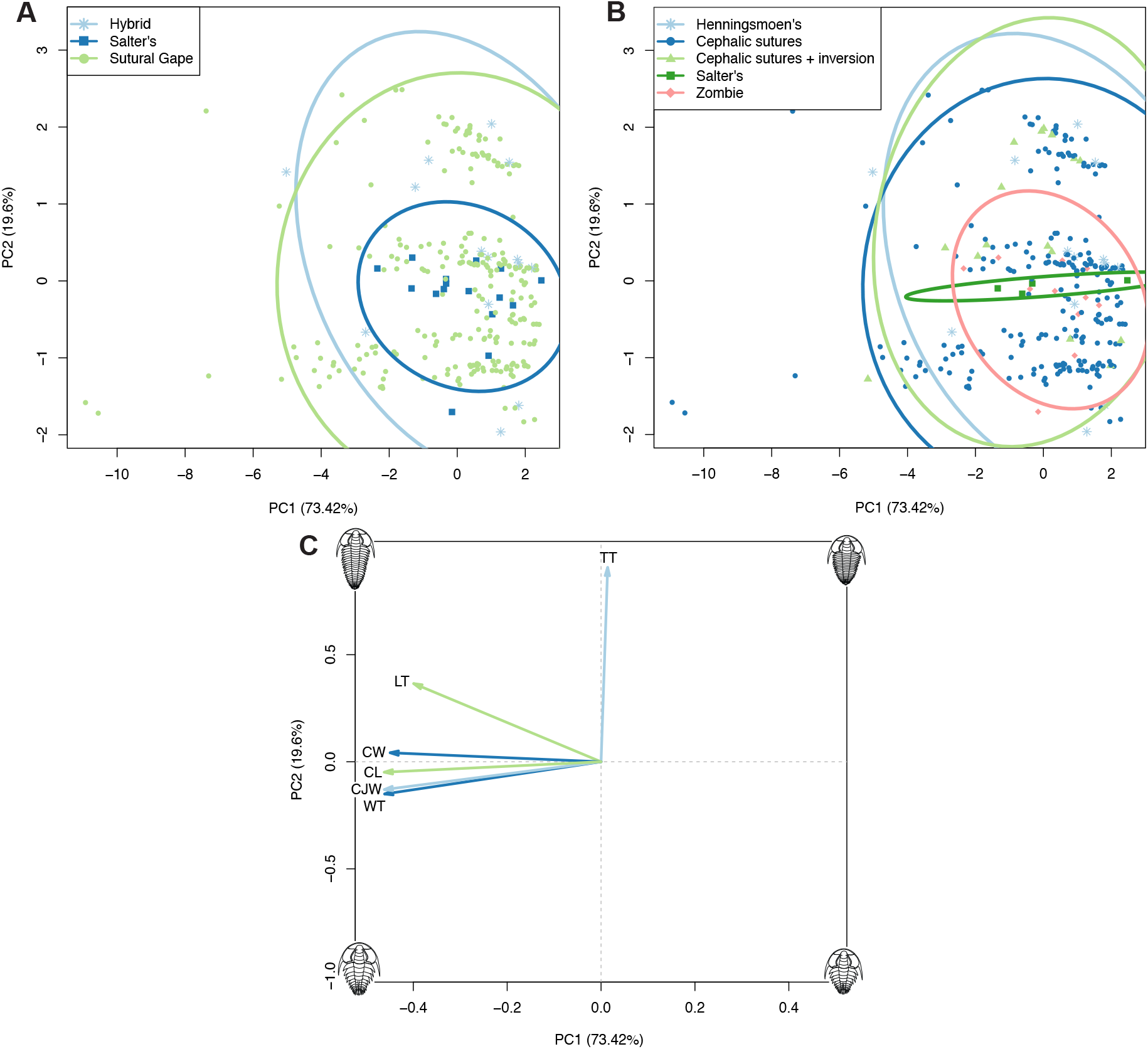
Principal Components Analysis plots with specimens grouped by moulting modes (A) or generalised moulting configurations (B) (see legends). Plot C shows the contributions of the dataset variables to the morphospace in A and B; directionality of each arrow represents how it varies across the morphospace, and length of each arrow represents its comparable impact on the morphospace. Abbreviations for C: TT, thoracic tergite number; LT, thoracic length; WT, thoracic width; CW, cranidial width; CL, cranidial length; CJW, cephalothoracic joint width.

**FIGURE 3.**
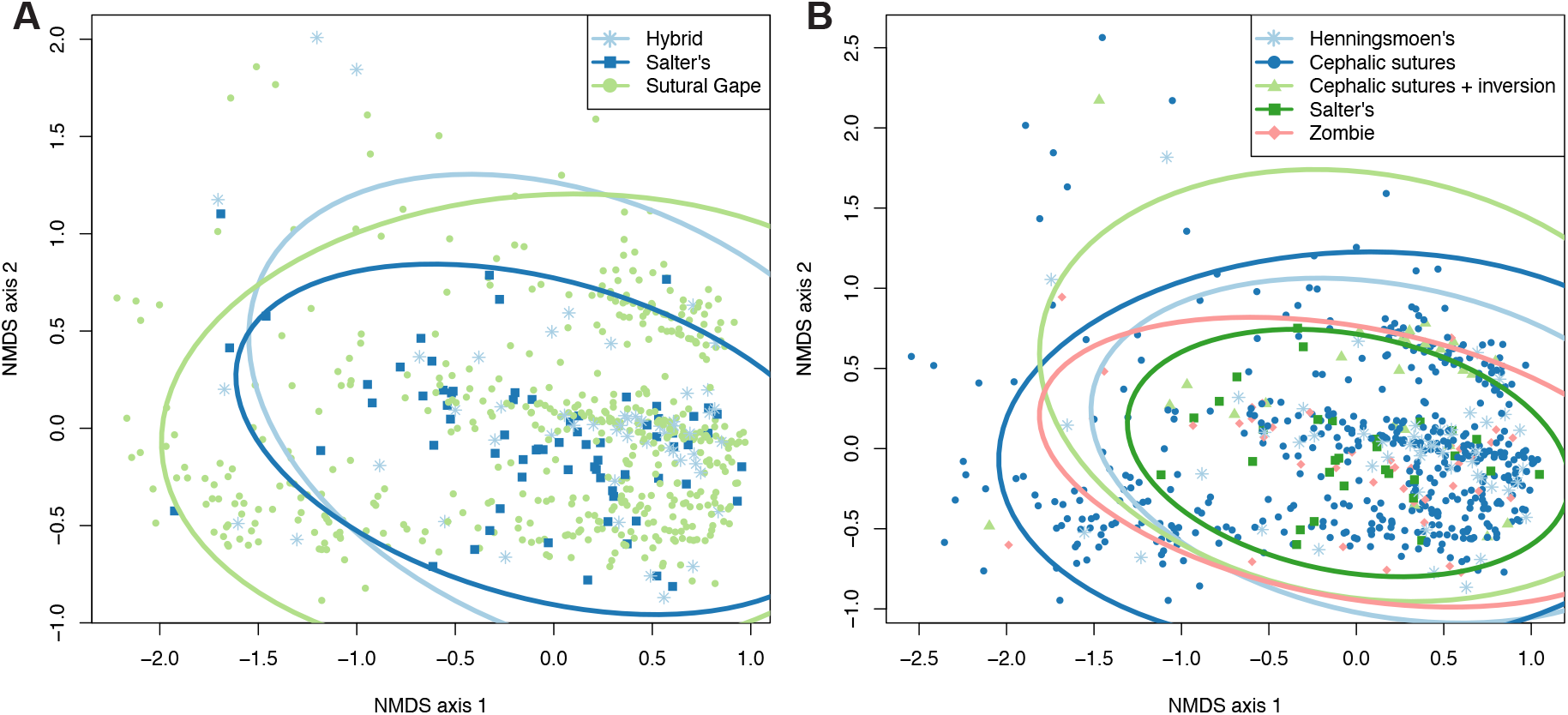
NMDS analysis plots with specimens grouped by moulting modes (A) or generalised moulting configurations (B) (see legends).

The PCA plots do suggest that Salter’s mode of moulting (Figure 2A) and its corresponding configurations (Salter’s and Zombie configurations; Figure 2B) occupy a smaller region of morphospace than the other moulting groupings. This suggests a restricted range of morphometries for trilobites using Salter’s mode, particularly in their number of thoracic tergites (along the PC2 axis) and their thoracic length (split between PC1 and 2 axes), this latter likely as thoracic length is related to tergite number (Figure 2C). The Zombie configuration grouping showed similar variation, though Salter’s configuration was even more restricted in thoracic tergite number and length (Figure 2B). This smaller morphospace area occupation is not solely an artefact of the PCA sample sizes of these groupings, because other groupings in the plots have equivalent sample sizes. However, the NMDS plots, which include the total dataset sample, do not appear to support this result (Figure 3), with all groupings showing similar extents of variation in the area of morphospace occupied.

### Thoracic tergite number and length are significantly different for moulting configurations

The median number of thoracic tergites significantly differed between the moult configuration groups (Table 4), demonstrated by a significant Kruskal-Wallis chi-squared test (χ^2^ = 2.828, *p* = 0.00103). Moult specimens in the cephalic sutures + inversion group are responsible for this significant result, as evidenced by post-hoc Wilcoxon pairwise tests that were significant for this group when paired with all other configuration groupings (Henningsmoen’s *p* = 0.00690, cephalic sutures *p* = 0.00077, Salter’s *p* = 0.00188, Zombie *p* = 0.00053; no other pairings significant). The cephalic sutures + inversion group had a higher thoracic tergite count, with a mean of 15 and median of 17, compared to a mean of 12 for all other groupings (and 10 for Zombie configuration; Table 4, Figure 4B). However, there was no significant thoracic tergite number difference when comparing moulting mode groups rather than configurations (Kruskal-Wallis: χ^2^ = 18.412, *p* = 0.243). Box plots of thoracic tergite number do show three major areas of clustering for the Sutural Gape mode, with the most extreme slightly skewing the results towards a high number of tergites (Figure 4A), but this is not reflected in the group averages. Sutural Gape mode specimens had a mean average of 12 thoracic tergites and Salter’s mode 11 tergites, with their medians both being 11 tergites, though the median for the hybrid mode was slightly higher at 13 tergites (Table 3, Figure 4A). The extreme positive cluster seen for the Sutural Gape mode in the boxplot (Figure 4A) seems more driven by the high tergite number for the cephalic sutures + inversion group, as the cephalic sutures configuration group has a confidence range with a lower top-end value (Figure 4B) compared to the Sutural Gape mode (Figure 4A). These findings all support that the significant results suggesting differences in thoracic tergite number are entirely due to the cephalic sutures + inversion configuration group, rather than a broad signal in the dataset. Although, tergite number box plots (Figure 4) may indicate a porous upper limit to the number of thoracic tergites for trilobites showing Salter’s and the hybrid modes of moulting (and their constituent configurations), as the numbers of specimens showing more than about 13 thoracic tergites for these groupings are extremely small.

**FIGURE 4.**
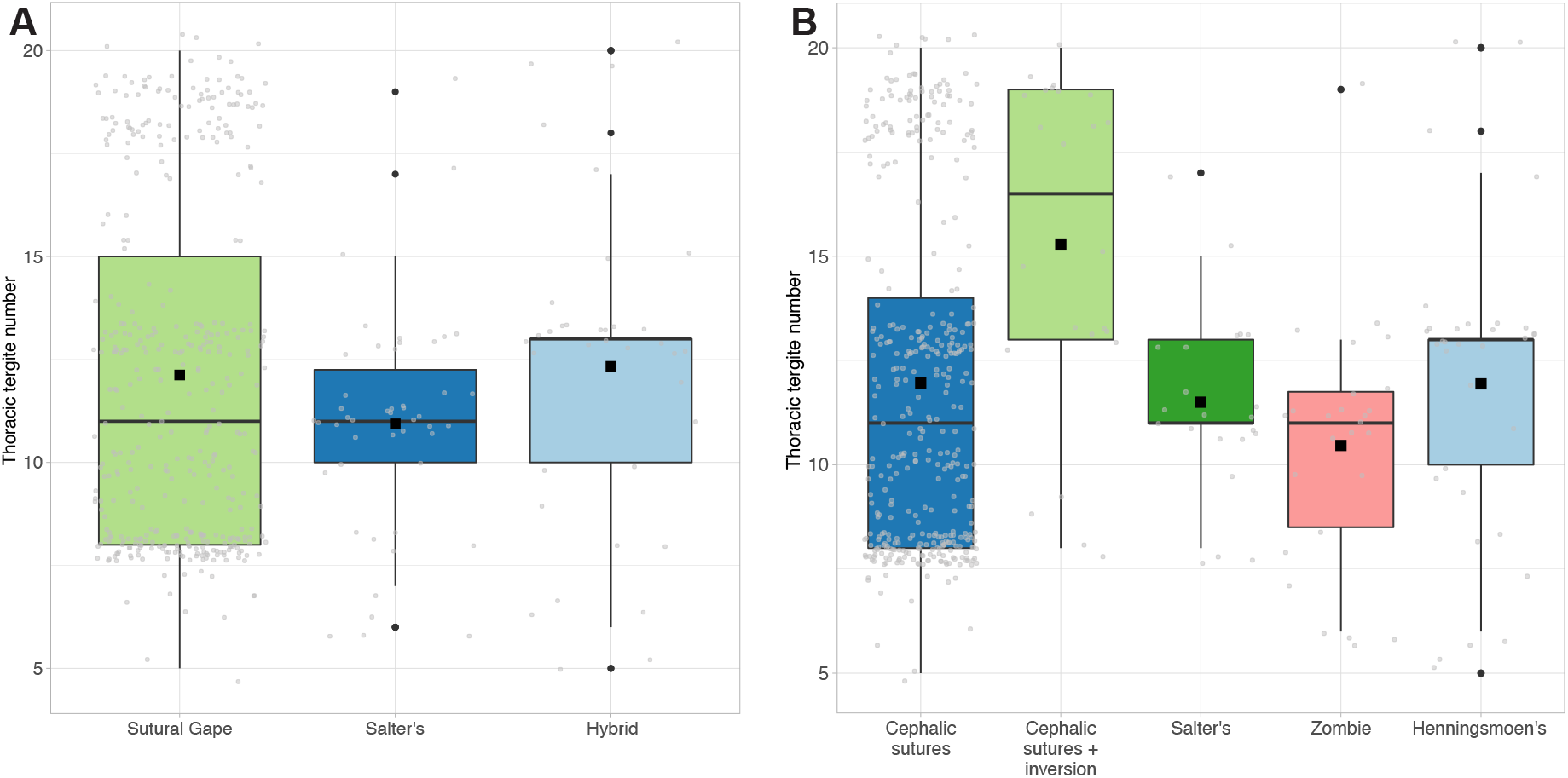
Box plots displaying mean (black square) and median (horizontal black line) average tergite number for each mode of moulting group (A) and each generalised moult configuration (B). Grey jitter shows the plotted points comprising the boxes, and black points show suggested outliers.

**TABLE 3.** Descriptive statistics for the dataset, displaying the mean, standard deviation, and median values for each morphometry variable, grouped by mode of moulting. The hybrid mode of moulting represents moult specimens in Henningsmoen’s configuration, with both the cephalic moulting sutures and the cephalothoracic joint opened.

| Mode of moulting | Morphometry variable | Mean (mm) | Standard deviation | Median (mm) |
| --- | --- | --- | --- | --- |
| Sutural Gape | thoracic tergite # | 12 | 4.2 | 11 |
|  | thoracic width | 22.4 | 14.8 | 17.4 |
|  | thoracic length | 18.1 | 12.1 | 15.7 |
|  | cephalothoracic joint width | 21.3 | 13.8 | 16.9 |
|  | anterior cranidium width | 13.4 | 9.8 | 10.8 |
|  | cranidium length | 13.7 | 8.6 | 11.2 |
|  | pygidium width | 16.8 | 16.1 | 10.8 |
|  | pygidium length | 10.1 | 10.1 | 5.9 |
|  | total body length | 39.2 | 25.5 | 31.1 |
| Salter’s | thoracic tergite # | 11 | 2.6 | 11 |
|  | thoracic width | 22.8 | 11.3 | 21.2 |
|  | thoracic length | 20.2 | 10.8 | 18.1 |
|  | cephalothoracic joint width | 20.9 | 11.7 | 18.8 |
|  | anterior cranidium width | 11.7 | 4.9 | 10.6 |
|  | cranidium length | 13.0 | 6.8 | 11.6 |
|  | pygidium width | 17.4 | 10.5 | 15.8 |
|  | pygidium length | 10.1 | 7.7 | 7.5 |
|  | total body length | 40.2 | 22.6 | 35.0 |
| Hybrid | thoracic tergite # | 12 | 3.9 | 13 |
|  | thoracic width | 20.4 | 13.0 | 15.7 |
|  | thoracic length | 19.9 | 17.7 | 14.0 |
|  | cephalothoracic joint width | 22.8 | 31.1 | 14.9 |
|  | anterior cranidium width | 11.2 | 7.1 | 8.7 |
|  | cranidium length | 11.7 | 7.7 | 9.8 |
|  | pygidium width | 13.3 | 11.8 | 9.0 |
|  | pygidium length | 7.59 | 7.2 | 5.6 |
|  | total body length | 32.3 | 22.8 | 24.6 |

**TABLE 4.**
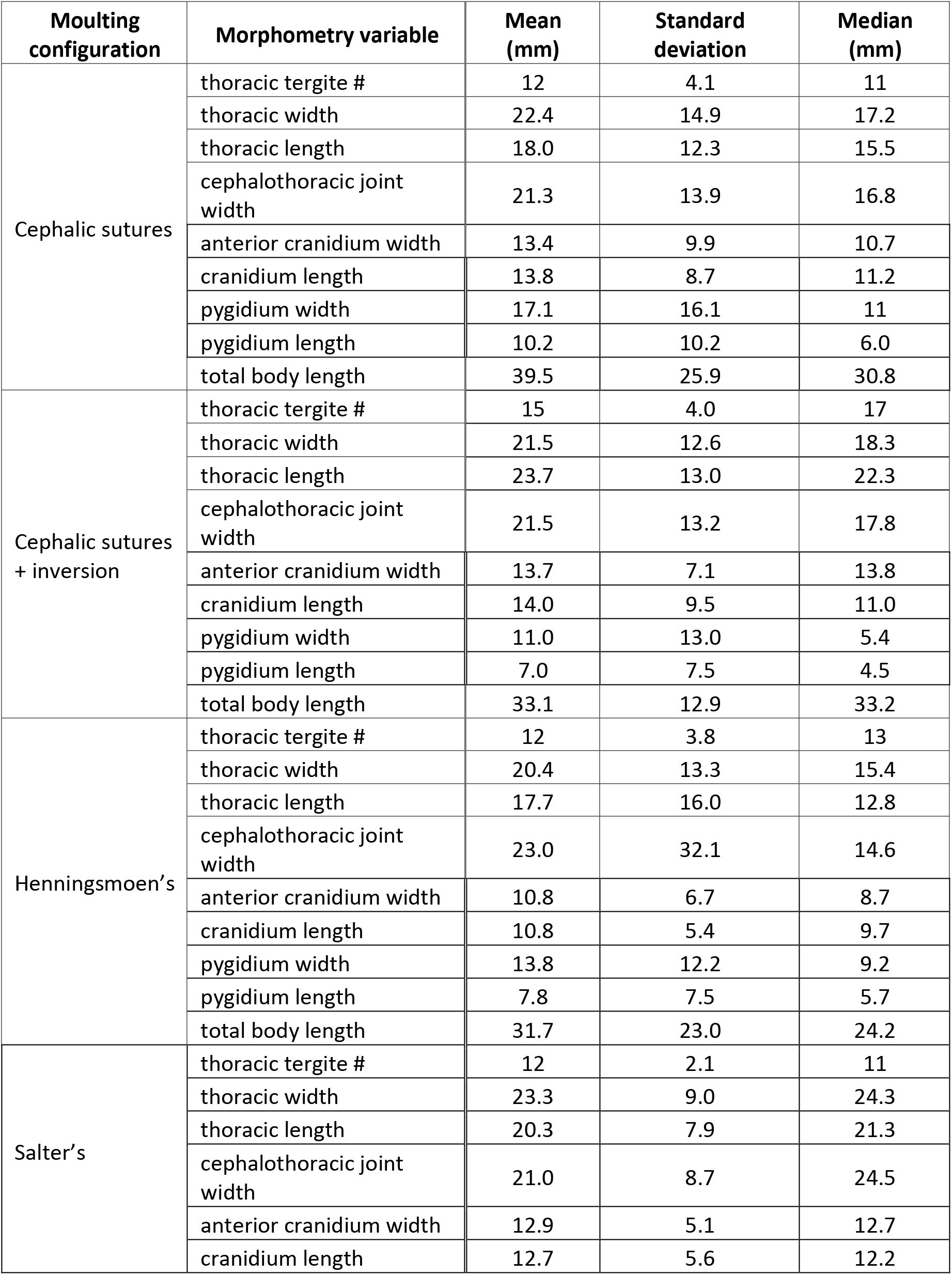
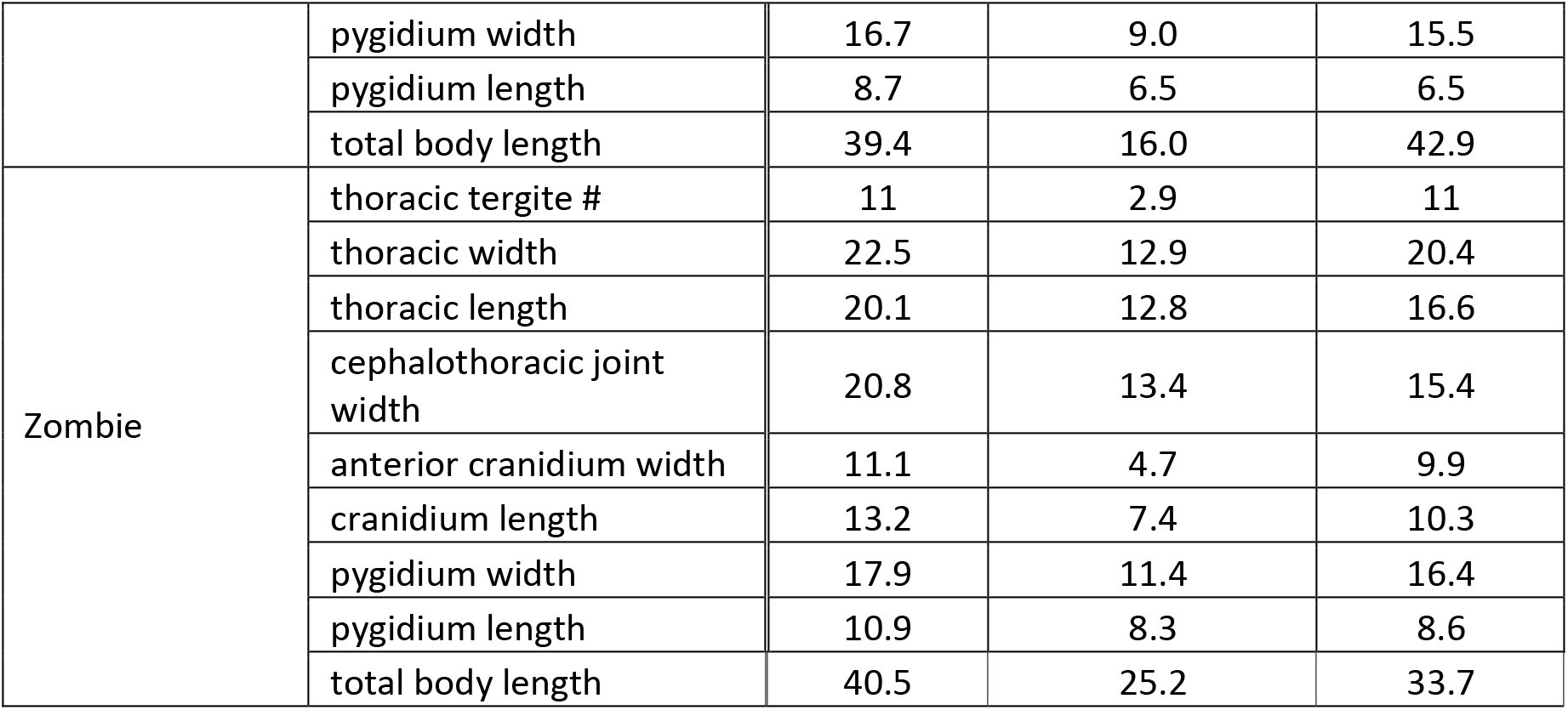
Descriptive statistics for the dataset, displaying the mean, standard deviation, and median values for each morphometry variable, grouped by moulting configuration.

As for thoracic tergite number, median thoracic length was significantly different between moult configuration groupings (Kruskal-Wallis: χ^2^ = 12.137, *p* = 0.0164), but not between moulting modes (χ^2^ = 2.845, *p* = 0.241). This again results from the cephalic sutures + inversion configuration group (Wilcoxon, *p* = 0.043 with cephalic sutures and Henningsmoen’s configurations; no other pairings significant), with the inversion group having a thorax with a mean length of 23.7 mm compared to 17.7 mm and 18.0 mm for the Henningsmoen’s and cephalic sutures groups, respectively (with the other two configurations having means of 20.3 and 20.1 mm). The similar results for the thoracic tergite number and length variables can be explained by the presumable co-dependence of the two, with a higher number of thoracic tergites generally associated with a longer thorax, and vice versa. Moulting mode and configuration were also determined to have a significant impact on thorax length variance based on a MANCOVA test (*p* = 0.0354 for moulting modes, *p* = 0.0210 for configurations).

### Pygidium width is significantly different for moulting configurations

Median pygidium width was also found to be significantly different between both moulting mode groups (Kruskal-Wallis: χ^2^ = 7.849, *p* = 0.0198) and configuration groups (χ^2^ = 12.703, *p* = 0.0128). For the moulting mode groupings this appears to result from the pairings of the Salter’s mode group (Wilcoxon: Sutural Gape and Hybrid both *p* = 0.0260), with Salter’s mode having a mean pygidium width of 17.4 mm compared to 16.8 mm for Sutural Gape and 13.3 mm for the hybrid mode (Table 4). When interrogating the configuration grouping result, only the cephalic sutures + inversion group was found to have a significantly different pygidium width to other configurations (Wilcoxon: Salter’s *p* = 0.0430, Zombie *p* = 0.00560; no other pairings significant). The inversion group had a narrower pygidium than these groups, at a mean of 11.0 mm compared to 16.7 mm and 17.9 mm respectively for the Salter’s and Zombie configuration groups. Additionally, both moulting mode and configuration were suggested to have a significant impact on pygidium width variance based on MANCOVA tests (*p* = 0.0109 for moulting modes, *p* = 0.0361 for configurations). This significant result is made clear by a scatterplot based on the MANCOVA model, with the linear regression lines clearly differing for the Sutural Gape and Salter’s modes of moulting (Figure 5F).

**FIGURE 5.**
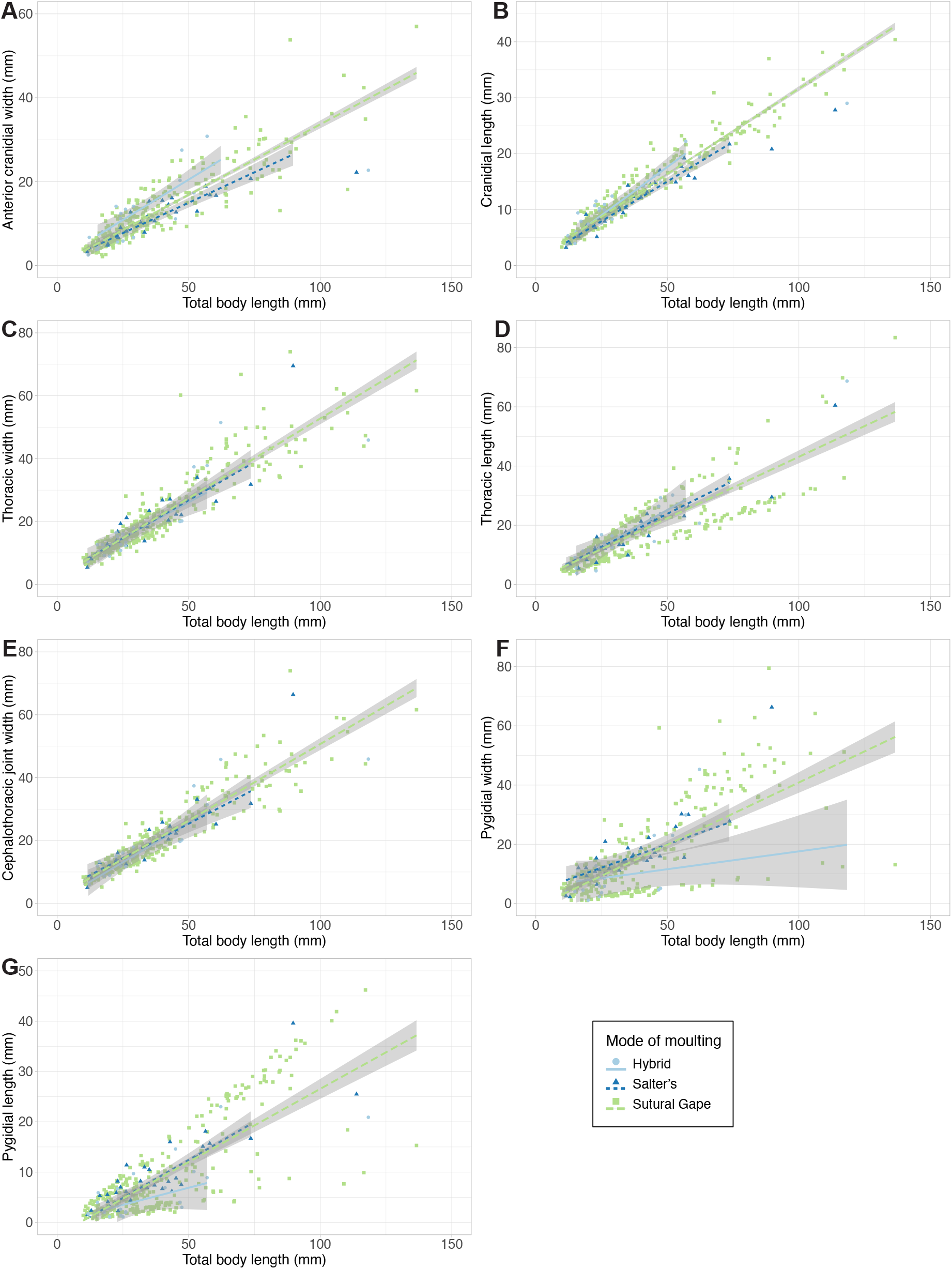
Scatterplots of the total dataset, for each morphometry variable. Data is grouped by mode of moulting (see legend). Linear regression lines represent the MANCOVA analysis models (see text).

### Moulting is unrelated to most other morphometric variables

The descriptive statistics suggest potential differences between the moulting groups for other morphometric variables (body length, pygidium length, cephalothoracic joint width, thoracic width, anterior cranidium width, cranidium length). For example, the median averages suggest specimens showing Salter’s mode of moulting were generally larger, and those showing the hybrid mode were generally smaller (Table 3). However, these remaining morphometric variables do not show statistically significant differences between moulting mode or configuration groups, based on Kruskal-Wallis tests. Further, MANCOVA tests indicate that both moulting mode and configuration had no significant impact on these morphometric variables, suggesting that these are unrelated to moulting behaviour. Scatterplots based on the MANCOVA models exemplify this lack of significant difference between the moulting groups for these variables (Figure 5A-C, E, H). These plots generally show a single linear positive correlation, indicating these body proportions increase in size linearly with body length, rather than having multimodal distributions or the moulting groups showing differing slopes.

### Multimodal distribution of results?

Despite the lack of clear difference in morphospace occupation of the different moulting groups, the PCA plots (Figure 2) suggest a potential non-unimodal distribution of the data. There are three distinct clusters of points observable in the PCA morphospace, separated primarily along PC2, and which appear to cross the moulting groupings rather than being related to moulting behaviour (see particularly Figure 2B). The NMDS plots show the same three clusters of points, again apparently unrelated to moulting group (see particularly Figure 3B). This clustering indicates a trimodal distribution (three overlapping unimodal distributions) in the total dataset. Additionally, at least two clusters of points can be seen in several of the linear regression scatterplots, including for at least thoracic length (Figure 5D), pygidial length (Figure 5G), and pygidial width (Figure 5F); this also indicates bimodal or trimodal distributions for some of the morphometric measurements. Finally, similar data clustering can be observed in the thoracic tergite number boxplots (Figure 4), particularly for the Sutural Gape mode and cephalic sutures configuration, though the data also suggest this pattern may hold for the other moulting groupings if given greater sampling.

Further exploration of distribution patterns in the dataset through interpretation of histograms suggests a right-skewed unimodal distribution for all continuous morphometric measurements, and only a potential multimodal distribution for thoracic tergite number (Figure 6C). EM algorithm analysis (Benaglia et al., 2009) supports this result for all variables, with all except for thoracic tergite number appearing to best fit a unimodal right-skewed distribution (e.g., Figure 6A; log-likelihood at convergence for two and three distribution peaks = −731 to −1035). Pygidium length and width (and to an extent cranidium width and length) show a slightly better fit with a bimodal distribution (with two means predicted by the EM algorithm) than the other variables (e.g., Figure 6B). However, a right-skewed unimodal distribution remains the best fit for these variables, as is clear from their histograms (Figure 6B underlying histogram). For thoracic tergite number, the EM algorithm analysis suggests a trimodal distribution (three overlapping unimodal distributions, each with a different mean) best fits the data (Figure 6C; log-likehood = −567.809), with a bimodal distribution having a poorer fit (log-likehood = −637.807). A trimodal distribution for the tergite number variable fits the three data clusters described from the morphospace plots (Figures 2 and 3) and boxplots (Figure 4), suggesting this represents a true signal in the dataset. The three morphospace clusters appear to correspond with the three distribution peaks predicted by the EM algorithm, because the clusters are indeed separated along PC2 (representing mainly thoracic tergite number variation) rather than PC1 (most other variables) (see Figure 6D).

**FIGURE 6.**
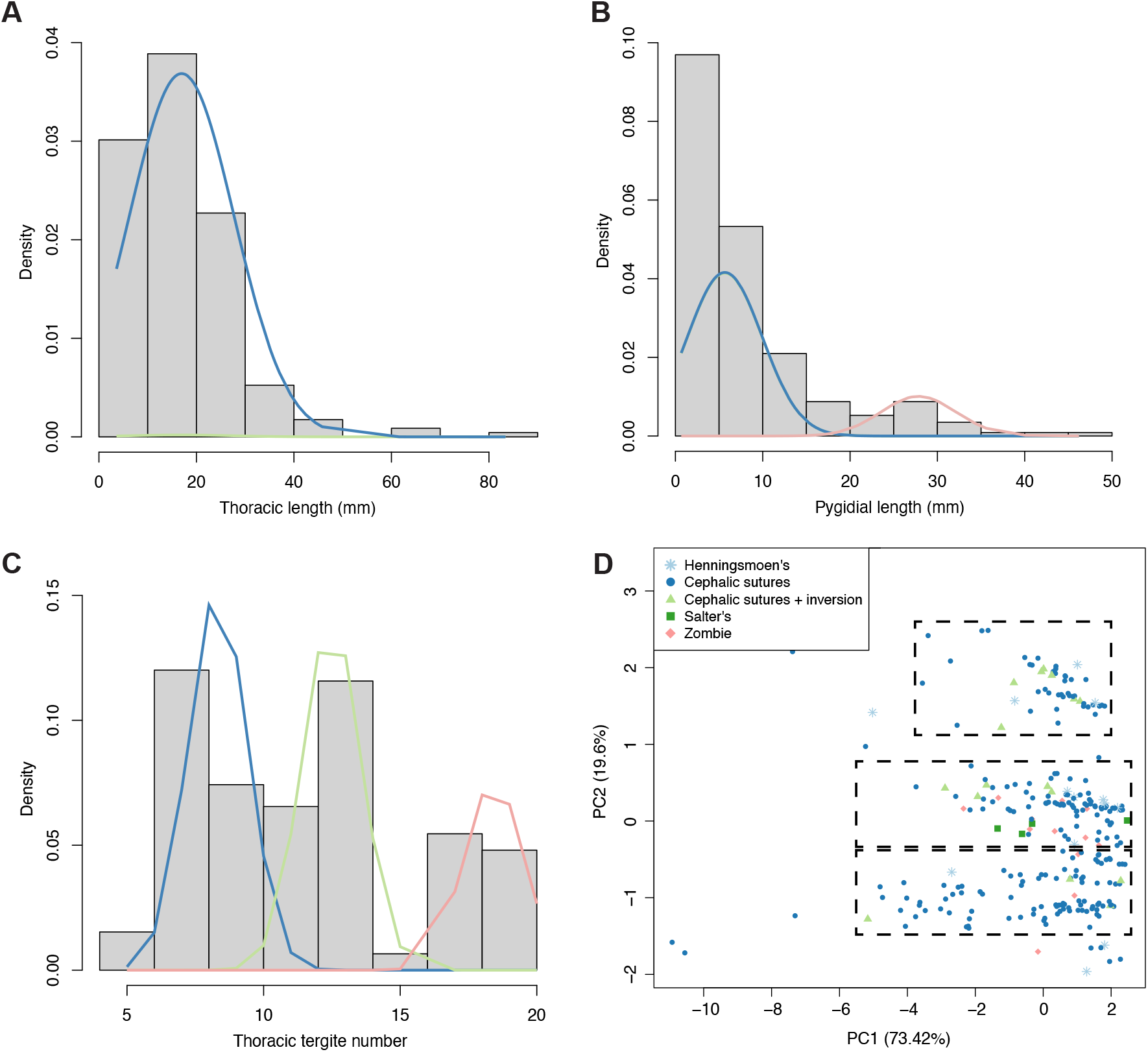
Histograms of notable variables within the dataset (using only complete data entries), with plotted EM algorithm-calculated distributions. A: thoracic length histogram, with two estimated distributions. B: pygidial length histogram, with three estimated distributions; two distributions (blue and green lines) are exactly overlapping, indicating they represent the same distribution. C: thoracic tergite number, with three estimated distributions. D: PCA plot, with specimens grouped by generalised moulting configuration (see legend) and ellipses removed (original plot in Figure 2B); dashed black boxes roughly demonstrate the three major data clusters apparent in the morphospace.

## DISCUSSION

Overall, the results of this morphometric study suggest that there is only a minor association between morphometry and moulting behaviour in trilobites. Certain results suggest a morphometric difference between individuals employing different modes of moulting, in particular the Salter’s mode group in PCA and NMDS analyses showing less variation than groups using the cephalic sutures (Figures 2, 3). This may suggest that the Sutural Gape mode of moulting is feasible over a wider range of body morphometries. Further, the morphospace plots and thoracic tergite number boxplots (Figure 4) suggest that the Sutural Gape mode may dominate at high tergite numbers. It may also be the case that trilobite species that habitually used Salter’s mode of moulting (particularly if resulting from fusion of the facial sutures) had a more constrained morphometry due to an evolutionary phenomenon like a phylogenetic bottleneck, as many of these species, though not all, belong to the same suborder (Phacopina; Crônier, 2013; Drage, 2019a). However, the NMDS analyses show the moulting groups to be more similar in their extents of variation, so the PCA results may be partially related to the necessary exclusion of incomplete specimen data in these analyses.

Most morphometric variables in the dataset were unrelated to moulting. However, means and variances of pygidium width did appear to significantly differ between the moulting behaviour groupings. Pygidium width significance may be related to the major pygidial forms described for trilobites (see Whittington et al., 1997; Hughes, 2007), and these different forms might also explain the apparent multimodal distributions observed in scatterplots (Figure 5F and G). Isopygous and macropygous pygidia tend to be broader than the thorax, and therefore may be more difficult to extract through potentially smaller exuvial gapes created by the cephalic sutures during moulting compared to micropygous pygidia. Indeed, pygidia were found to be significantly broader in specimens showing Salter’s mode of moulting compared to the Sutural Gape and hybrid modes (both of which use the cephalic sutures; Table 3). At a moulting configuration level, the cephalic sutures + inversion group had a significantly narrower pygidium in comparison to the broader pygidium seen in the Salter’s and Zombie configuration groupings (Table 4). Histograms of separate morphometric measurements suggest that neither pygidium measurements are multimodally distributed with peaks corresponding to macro-micropygous forms, although a potential minor second distribution is interpretable from the EM algorithm analysis (Figure 6B). Based on these results, this apparent pygidium size sorting in the dataset may be partially related to moulting behaviour differences, rather than just reflecting the different morphological forms known in trilobites.

Thoracic length means and variances were also significantly different between the moulting groups in the dataset. Moult specimens with only open cephalic sutures appeared to have an overall shorter thorax than those using the cephalothoracic joint during moulting (Table 3), which may be for mechanistic reasons such as a longer thorax being easier to extract through a joint at the thorax than a potentially thinner gap at the anterior of the cephalon. Though a stronger trend would be required for this interpretation to be convincing, and the Kruskal-Wallis comparisons suggest it is only the cephalic sutures + inversion configuration specimens that significantly differ in thorax length. There was also a significant association between moulting configuration and thoracic tergite number, caused by the cephalic sutures + inversion configuration specimens having a higher number of tergites (Figure 4). Brandt (2002) suggested that an overall trend to decreased thoracic tergite numbers over trilobite evolutionary history (Hughes et al., 2001) might be related to fewer tergites being less risky for moulting. The minor association between moulting behaviour and tergite number reported here does not seem to bear this out, with the cephalic sutures + inversion group unlikely to be much riskier than the highly common cephalic sutures group. The results here may be due to the reasonably small sample size for the cephalic sutures + inversion group (25 specimens), though the otherwise lack of association between tergite number and moulting behaviour is probably not sampling-related due to the generally large sample sizes. Given this absence of broad signal regarding thoracic length and tergite number in the dataset, it is reasonable to suggest this result may be related only to mechanical mechanisms such as articulation strength. This result is the opposite to that found by Drage et al. (in press), in which *Estaingia bilobata* moults with longer bodies were often preserved with only cephalic sutures open, potentially suggesting better developed articulations with increasing size reduced the incidence of Salter’s mode of moulting. This contrarian result suggests that this thoracic morphometric trend cannot be applied across this diverse group, and perhaps reflects lower taxonomic level biomechanical differences.

The apparent non-unimodal distributions present for some variables within the dataset raise additional interesting considerations about potential morphological constraints in trilobites. A unimodal skewed (rather than normal) distribution, which we see for most morphometric variables in this dataset (e.g., Figure 6A), is common and can be informative for continuous measurements of biological systems (e.g., Church et al., 2019), particularly for body size (Kozłowski and Gawelczyk, 2002). However, it is interesting that clustering in the linear regression scatterplots (Figure 5) and the EM algorithm results (Figure 6B) suggest the potential for a bimodal distribution in pygidium measurements. As noted above, these clusters, or distribution peaks, might represent different morphometric trajectories for groups with highly differing morphologies or ontogenetic pathways. The clear multimodal distribution for thoracic tergite number (Figure 6C), which explains the multiple clusters seen in morphospace plots (Figures 2 and 3), perhaps suggests that trilobites tend to congregate at certain numbers of thoracic tergites with this dataset suggesting more trilobites with c. 8, 12, and 18 thoracic tergites than between these (Figure 6). This is interesting because of the highly complex and varied segmentation patterns occurring within the Trilobita (e.g., see Hughes, 2003). Apparent stability of these thoracic tergite numbers could be explained by many factors affecting broad-scale evolution that are difficult to unpick, including morphological, behavioural, and ecological factors. Hughes et al. (2001) discussed convergence on stable thoracic tergite numbers as a broad-scale evolutionary pattern visible in trilobites. Tergite number stability is likely only tangentially related to moulting behaviour because a broad-scale signal is not apparent in the dataset (only for cephalic sutures + inversion configuration specimens). Peaks in thoracic tergite number, such as that seen in this dataset, could also be explained by evolutionary bottlenecks, in which tergite number is more controlled by phylogenetic descent than by adaptation. Stabilisation of tergite number in trilobites could thereby result from a constraint on this number in early representatives of the group, and through time sufficient morphological adaptation took place around this tergite number to cause alteration of this characteristic to be evolutionarily unsuccessful. Indeed, Hughes (2003) noted that early-diverging trilobite groups displayed high variation in thoracic tergite number, and derived clades showed more stable numbers of tergites, with higher taxonomic ranks in particular showing more stable tergite numbers later in trilobite evolutionary history (Hughes et al., 2001).

Moult configurations with exoskeletal inversion, both of cephalic sclerites and the complete cephalon (Salter’s configuration), were generally uncommon in the dataset, potentially suggesting these were a rarer occurrence in Trilobita than may be highlighted in certain species (Whittington, 1990). Inversion has been suggested to occur through dorsal flexure or partial enrolment of the individual during moulting (Whittington, 1990; McNamara and Rudkin, 1984; Daley and Drage, 2016; Drage, 2019a,b), for example in *Estaingia bilobata* (Drage et al., 2018a), and so its rarity here suggests many trilobites may have remained unflexed during moulting. Although, changes in moulting behaviour through ontogeny described by Wang et al. (2021) may suggest that certain trilobite species did regularly employ flexure for moulting but that this was linked to their development. Alternatively, this may reflect a description bias of trilobite moult specimens. Interesting or unusual moult specimens may, in general, be preferentially figured within manuscripts, but when assessing the bulk of historical museum specimens these may be less common in number than expected.

## CONCLUSIONS

This work presents the first large cross-group quantitative analysis exploring a potential link between trilobite moulting behaviour and morphology. The dataset suggests some minor differences in morphometry between groups of specimens displaying different modes of moulting and preserving different moult configurations. In particular, the apparent presence of a longer thorax with more articulations and a narrower pygidium in specimens showing open cephalic sutures + inversion, and potentially less variation in the morphology of specimens using Salter’s mode of moulting. However, little association between most of the morphometry variables investigated and any other moulting grouping was detected. Further, although the sample size for the cephalic sutures + inversion configuration group was sufficient to be included in this study (at 25 specimens), more specimens (ideally approximately 100) would be required for the results to be considered conclusive, though this would be difficult due to the configuration’s apparent rarity. Lastly, if there was a clear association, even minor, between morphometry and moulting behaviour in trilobites, we would not expect to see significant results with only one, reasonably rare, grouping of moult configurations. To be confident of any real biological association, we would require some suggestion of differing morphometry between the main modes of moulting; Sutural Gape and Salter’s modes.

The lack of clear morphometric association with other moulting groupings in the dataset is itself interesting, because we would expect morphology and key behaviours like moulting to interact and invoke changes in each other, particularly over long evolutionary timescales (Henningsmoen, 1975; Brandt, 2002). It raises the question of whether moulting behaviour variability in trilobites can be considered related to morphological aspects outside of the obvious existence or loss of physical moulting mechanisms, such as the facial sutures (Whittington, 1990; Drage, 2019a). There may be a minor association, as suggested by this morphometric study, however other factors influencing morphology seem to have played larger roles in determining moulting behaviour variability. For example, it is possible that, rather than being directly related to morphometry, moulting behaviour interacted with morphological complexity, as Brandt (2002) suggested, such as the number of exoskeleton articulations and spinosity (the latter of which Whittington, 1990, suggested may be important as leverage during moulting). The thoracic tergite count results presented here do not necessarily bear this out, though broad-scale trends in trilobite spinosity deserve further exploration and quantitative study of other complexity measures across groups could be consequently informative. Otherwise, this lack of strong morphometric signal suggests other aspects of trilobite evolutionary history that may have influenced moulting variability deserve attention, for example, phylogenetic relationships, and that greater in-depth study on trilobites is required to understand the evolutionary impacts of their moulting behaviour flexibility.

## ACKNOWLEDGEMENTS

Data collection was performed during a Natural Environment Research Council studentship 2014-2018 (NE/L002612/1), and analyses and writing under funding by the Swiss National Science Foundation Sinergia grant (198691). I wish to gratefully acknowledge the support, advice, and discussions of A.C. Daley and S. Pates over the full extent of this project, and their help in improving this manuscript. C. Mellish (NHMUK), J.O. Ebbestad (PMU), M.A. Binnie (SAM), K. Riddington (BIRUG), and E. Howlett (OUMNH) provided access to and support with museum collections, and data collection would have been impossible without them.

**SUPPLEMENTARY 1.** All raw data used for the analyses presented here are made freely available with this paper (Drage_Supplementarydata.xlsx).

